# Retrosplenial inputs drive diverse visual representations in the medial entorhinal cortex

**DOI:** 10.1101/2023.10.03.560642

**Authors:** Olivier Dubanet, Michael J. Higley

## Abstract

The ability of rodents to use visual cues for successful navigation and goal-directed behavior has been long appreciated, although the neural mechanisms supporting sensory representations in navigational circuits are largely unknown. Navigation is fundamentally dependent on the hippocampus and closely connected entorhinal cortex, whose neurons exhibit characteristic firing patterns corresponding to the animal’s location. The medial entorhinal cortex (MEC) receives direct projections from sensory areas in the neocortex, suggesting the ability to encode sensory information. To examine this possibility, we performed high-density recordings of MEC neurons in awake, head-fixed mice presented with simple visual stimuli and assessed the dynamics of sensory-evoked activity. We found a large fraction of neurons exhibited robust responses to visual input that shaped activity relative to ongoing network dynamics. Visually responsive cells could be separated into subgroups based on functional and molecular properties within deep layers of the dorsal MEC, suggesting diverse populations within the MEC contribute to sensory encoding. We then showed that optogenetic suppression of retrosplenial cortex afferents within the MEC strongly reduced visual responses. Overall, our results demonstrate the the MEC can encode simple visual cues in the environment that can contribute to neural representations of location necessary for accurate navigation.

## Main Text

Spatial navigation is fundamentally dependent on the hippocampus and closely connected entorhinal cortex ^1,2^. However, navigation also requires the integration of sensory cues and internally generated motor signals ^3^. The combination of these various streams of information are thought to give rise to canonical patterns of spiking that correspond to environmental locations and features. For example, place cells in hippocampal CA1 exhibit action potential firing that maps on to specific places ^4^ and grid cells in the MEC fire at vertices of a lattice spanning the local environment ^5^. However, MEC neurons are also responsive to behavioral and environmental features including borders, task-relevant timing, and speed of locomotion ^6-8^, and there is growing appreciation that sensory cues can influence MEC activity ^9^.

The ability of rodents to use visual cues for successful navigation and goal-directed behavior has been long appreciated ^4,9-11^. Visual information can anchor place and grid cells during navigation ^5,12,13^, and loss of visual inputs or decoupling of the relationship between visual flow and self-motion can disrupt the activity of hippocampal and MEC neurons ^11,12,14^. Moreover, “cue” cells in the MEC exhibit repeatable firing fields near salient visual features ^9,11^. Computational work suggests the importance of visual information as a mechanism for error correction during path integration ^15,16^. In more recent years, investigators have used virtual reality tasks in which the rodent navigates an environment consisting only of visual cues, finding normal emergence of grid cell dynamics ^9^. One possibility is that neurons in the MEC inherit complex sensory representations from upstream areas like the retrosplenial cortex (RSP), a neocortical area that also appears to encode relational properties of visual landmarks ^17^. In contrast, the MEC may directly encode simple visual information, allowing local entorhinal and hippocampal circuits to integrate such cues with more complex behaviorally-relevant signals.

Traditionally, multimodal and highly processed unimodal sensory inputs arising from the neocortex are thought to target superficial layers in the MEC, whose cells relay those signals to the hippocampus ^18-20^. However, neurons in occipital neocortical regions including primary and secondary visual cortex as well as RSP also innervate deeper layers ^19-21^, and optogenetic activation of RSP fibers in the MEC *ex vivo* can evoke excitatory postsynaptic potentials in layer 5 neurons ^21^. Overall, the identity of visually responsive cells in the MEC and their sensitivity to specific visual afferents is unknown.

Here, we performed high-density recordings of MEC neurons in awake, head-fixed mice presented with simple visual stimuli and assessed the dynamics of sensory-evoked activity. Our results demonstrate that ∼40% of MEC neurons exhibit robust, time-locked activity in response to visual input. Visually responsive cells appear to comprise molecularly-distinct subpopulations in layers 3 and 5 that differ in their response profiles. Moreover, visual inputs strengthen the coupling of MEC output to ongoing network dynamics, suggesting a role for shaping circuit activity linked to behavior. Finally, we find that suppressing retrosplenial inputs to the MEC significantly disrupts visually driven activity, suggesting this pathway is a key mediator of visual representations in the hippocampus.

## Results

### MEC neurons are responsive to simple visual stimuli

To study sensory-evoked neural dynamics in the MEC, we recorded single unit and LFP activity in head-fixed, freely-running mice presented with small patches of contrast-modulated drifting gratings (Figure 1A, see Methods). Neuronal output was sensitive to changes in behavioral state, including fluctuations in running and pupil diameter as well as onsets of visual cues (Figure 1B). Indeed, we found that a surprisingly large fraction of MEC neurons was highly modulated by visual input (Figure 1C, see Methods. Most responsive cells (38% of all neurons) exhibited increased firing rates during the stimulus presentation, while a smaller proportion (4%) were suppressed (Figure 1C).

We next investigated the potential for single neurons to represent specific features of the visual stimuli in their signal-to-noise ratio (SNR), defined as (visual response)/(baseline firing rate). We found that, across animals, evoked activity increased with stimulus contrast (Figure 1D). Additionally, MEC visual responses were monotonically enhanced by increasing stimulus size (Supplemental Figure 1). However, responses were largely unaffected by the spatial frequency or orientation of the stimulus and were similar for both drifting and static (inverting) gratings (Supplemental Figure 1). We also found that visually evoked activity was somewhat greater for stimuli presented solely to the contralateral versus ipsilateral eye (Supplemental Figure 1). Overall, these results are consistent with a view of the MEC as an upper-level visual area, with high sensitivity and large receptive fields but low specificity for simple sensory cues.

**Figure 1.**
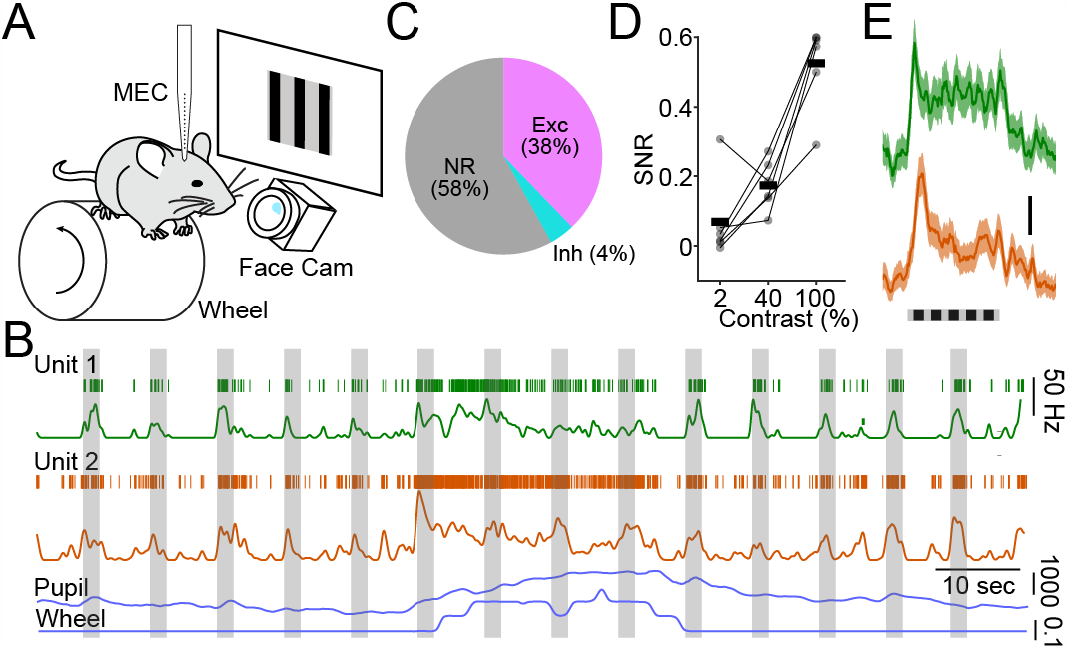
MEC neurons are responsive to simple visual stimuli. **A**, Schematic illustrating recording setup. **B**, Example recordings for two single units (green and orange) in the dorsal MEC. For each unit, upper trace shows raster plot of single spikes, lower trace shows instantaneous firing rate. Simultaneous pupil diameter and locomotion speed are shown at bottom (blue). **C**, Proportion of all recorded neurons exhibiting increased or suppressed firing during visual stimulation. **D**, Within-animal averages (n=6 mice) of visually-evoked SNR as a function of stimulus contrast. **E**, Average visual responses for the two units shown in (B).

### Heterogeneous visually-responsive subpopulations in the MEC

The MEC comprises a highly diverse population of neurons that vary as a function of both cortical layer and position along the dorsal-ventral axis ^18^. Therefore, we next sought to characterize the identity of MEC neurons responsive to visual inputs. First, we examined the presence and magnitude of visual responses as a function of recording location (determined by diI-labeling of the electrode). A greater percentage of cells were responsive and exhibited a significantly larger SNR for the dorsal versus ventral half of the MEC (p=0.011, linear mixed effects model, Figure 2A).

**Figure 2.**
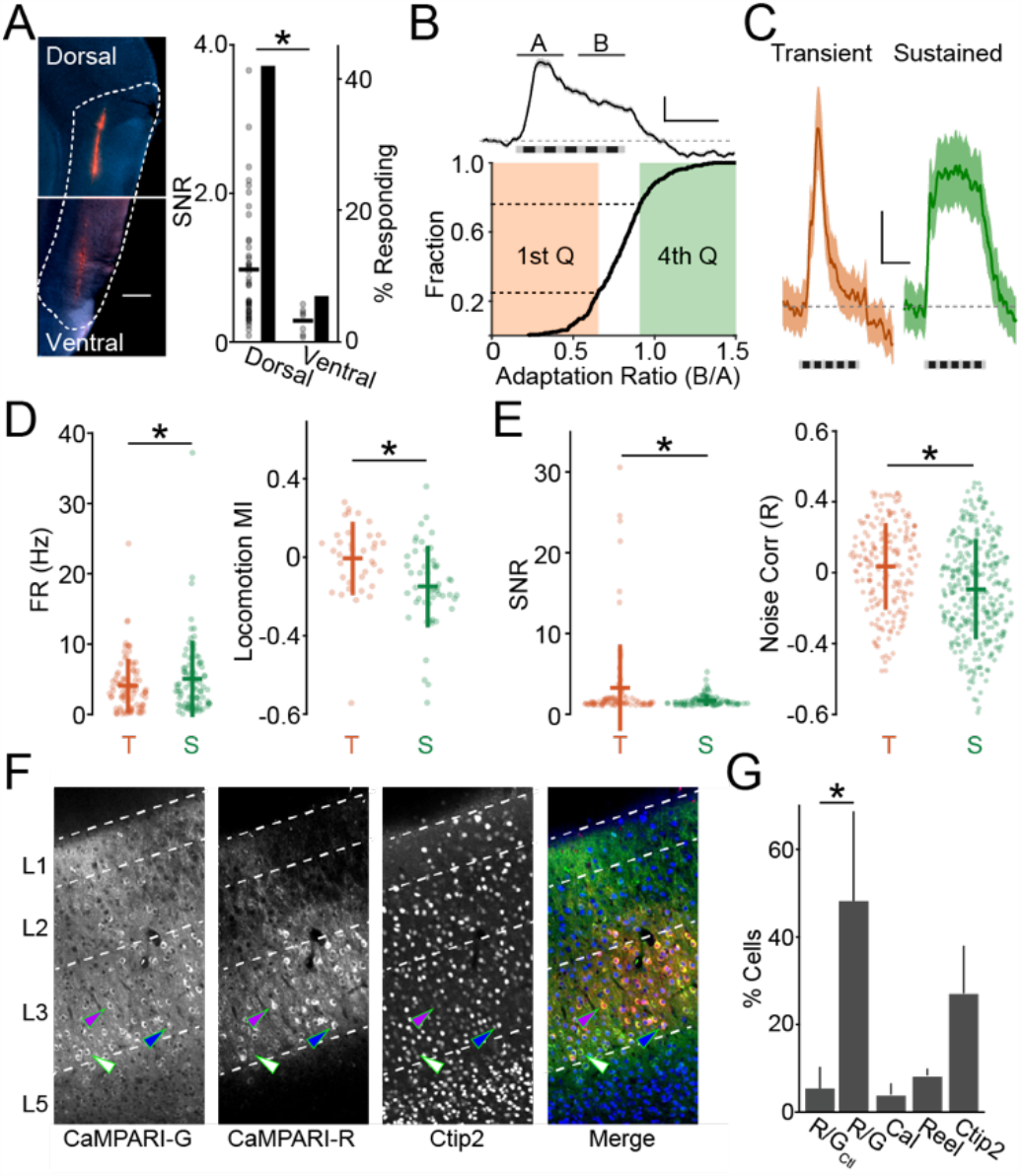
Visually responsive MEC neurons exhibit diverse functional and anatomical features. **A**, DiI-labeling of recording electrode track in dorsal MEC (left). Visual SNR (dots) and percentage of visually responsive cells (bars) for dorsal versus ventral MEC recording sites. **B**, Cumulative distribution plot of adaptation ratio (B/A, see example upper trace) for all visually responsive cells. Lower and upper quartiles indicated. **C**, Average responses across all cells in the lower (Transient) and upper (Sustained) quartiles for adaptation ratio, as in (B). **D**, Average spontaneous firing rates for transient versus sustained cells (left). Average locomotion Modulation Index for spontaneous activity for transient versus sustained cells (right). **E**, Average visual SNR for transient versus sustained cells (left). Average pairwise noise correlations for visual responses in transient versus sustained cells (right). **F**, Example images showing green-fluorescent CaMPARI2-labeled cells (1st panel), photoconverted red fluorescent cells (2nd panel), immuno-labeled Ctip2-positive cells (3rd panel), and merged image (4th panel). Individual non-converted cell (green arrow), converted Ctip2-positive cell (blue arrow), and converted Ctip2-negative cell (purple arrow) are indicated. **G**, Average percentage of cells indicated for photoconversion in control (no visual stimulation) animals, photoconversion after visual stimulation, and calretinin-, reelin-, or Ctip2-positive neurons. * indicates p<0.05 (see Main Text).

We confirmed this finding using fos-TRAP transgenic mice that allow fluorescent tdTomato-labeling of active neurons within a brief temporal window ^22^. We maintained fos-TRAP-expressing mice in the dark for five days. On the sixth day, mice were injected with hydroxy-tamoxifen, head-fixed on a running wheel, and presented with visual stimuli (drifting gratings) for 30 minutes. After 1 week, mice were perfused and td-tomato expression was quantified within the MEC. As expected from our recordings, red fluorescent cells were located primarily in the dorsal MEC, with the highest density of cells in layers 3 and 5 (Supplemental Figure 2).

Many visually responsive MEC neurons exhibited a transiently higher output at stimulus onset that decayed rapidly to a sustained elevated rate (Figure 1E). We quantified this tendency for single cells by calculating an adaptation ratio of firing during the second 500ms versus the first 500ms of the stimulus for all cells across all animals (Figure 2B-C, Supplemental Figure 3). Cells in the lower and upper quartiles were defined as transient and sustained, respectively, and investigated further to explore population differences in firing dynamics. These two categories did not differ in spike waveform from each other or from visually non-responsive cells (Supplemental Figure 3). However, sustained cells exhibited a modestly but significantly higher spontaneous firing rate (4.0±3.8 Hz versus 5.0±5.4Hz for transient versus sustained, n=354 cells in 33 mice, p=0.036, linear mixed effects model, Figure 2D) and were less sensitive to modulation of spontaneous activity by locomotion (Modulation index 0.19±0.18 versus 0.05±0.20 for transient versus sustained, n = 182 cells in 14 mice, p=0.0013, linear mixed effects model, Figure 2E). Surprisingly, MEC cells were not strongly modulated during changes in arousal measured by pupil diameter and the two groups did not differ (Supplemental Figure 3)

**Figure 3.**
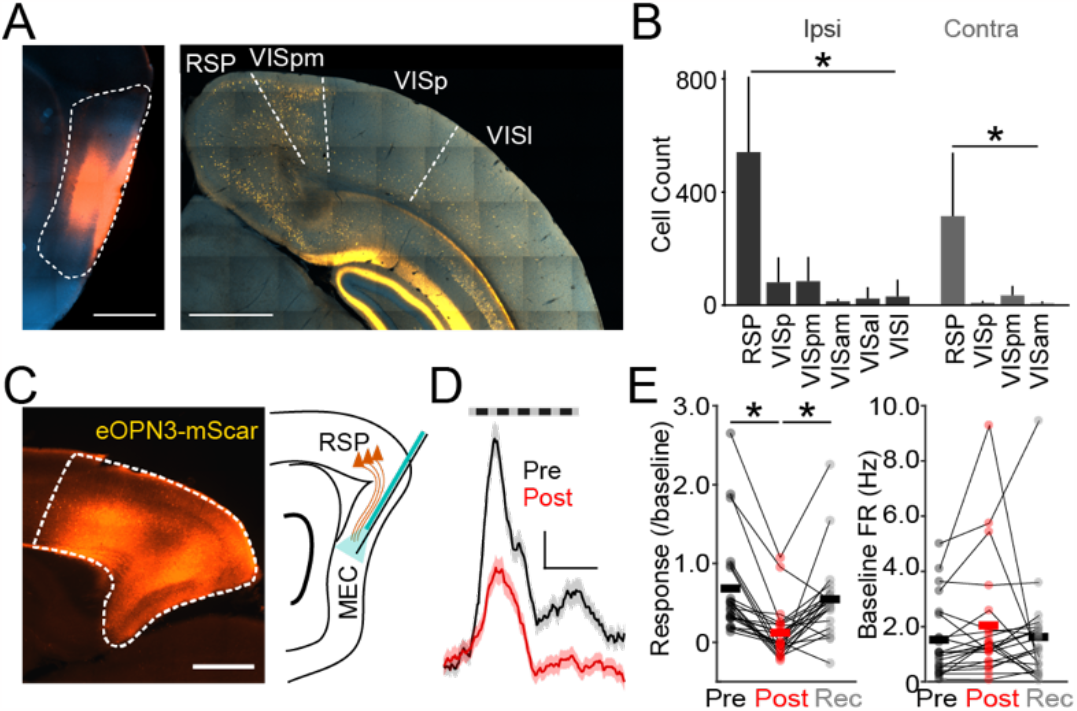
MEC afferents from RSP mediate visual responses. **A**, Example image showing expression of fluorescent CTB-Alexa Fluor-594 label in dorsal MEC (left). Example image of retrogradely labeled cells in ipsilateral neocortical visual areas (right). **B**, Average number of retrogradely labeled cells in different visual areas ipsilateral or contralateral to injection site. **C**, Example image showing expression of eOPN3-mScarlet in RSP (left) and schematic of simultaneous optogenetic activation and electrophysiological recording in the MEC. **D**, Average visual responses in MEC neurons before and after optogenetic suppression of RSP afferents. **E**, Average visual responses of MEC neurons relative to baseline before, immediately after, and 30 minutes after optogenetic suppression of RSP inputs (left). Average spontaneous firing rate of MEC neurons before, immediately after, and 30 minutes after optogenetic suppression of RSP inputs (right). * indicates p<0.05 (see Main Text).

We then analyzed visually evoked activity, finding that transient cells exhibited a significantly greater SNR (n=354 cells in 33 mice, p=0.0375, linear mixed effects model, Figure 2F). However, in contrast to spontaneous activity, visual responses were not altered by variation in the either locomotion or pupil diameter (Supplemental Figure S3). We also calculated pairwise noise correlations for simultaneously recorded, visually responsive MEC cells. This metric is thought to reflect the degree of either shared synaptic connectivity or common synaptic inputs ^23^(see Methods). Comparing within groups of cells, we found that noise correlations were significantly higher among transient versus sustained cells (n=354 cells in 33 mice, p=0.035, linear mixed effects model, Figure 2G), suggesting this group may comprise a distinct and more strongly interconnected subpopulation.

Within the MEC, theta oscillations (6-12Hz) in the local field potential (LFP) shape the timing of spike output from individual neurons and may link behavior to activity patterns of neuronal ensembles ^13^. We observed that visual stimulation significantly increased LFP power in this range (Supplemental Figure 3). In addition, visual input enhanced neuronal phase-locking to theta similarly for both transient and sustained cells (Supplemental Figure 3).

Our electrophysiological recordings suggest that visually responsive cells may comprise distinct subpopulations within the MEC. To further investigate the identity of these neurons, we expressed the activity-dependent fluorescent marker CaMPARI2 in the MEC (see Methods). We used an implanted fiber optic to deliver UV (405 nm) light to the MEC during presentation of visual stimuli identical to our recording experiments. Our results showed that CaMPARI2 expression was successfully targeted to deeper layers of the MEC (Figure 2F, Supplemental Figure 2). Moreover, approximately 45% of cells were photoconverted by delivery of visual stimulation (Figure 2G), similar to the proportion of visually responsive neurons by recording. Omission of visual stimulation resulted in only ∼5% of cells being photoconverted, suggesting red labeling is specific to visually responsive neurons. To determine the molecular signatures of visually responsive cells, we next immuno-stained CaMPARI2-labeled tissue for markers of known MEC subtypes, including calretinin, reelin, and Ctip2. Of these, less than 10% of photoconverted cells were positive for either calretinin or reelin, but ∼25% were positive for Ctip2 (Figure 2G, Supplemental Figure 2).

### Retrosplenial cortex drives visual responses in the MEC

We next investigated which inputs to the MEC might contribute to visual responses. We first labeled inputs to the dorsal MEC via injection of fluorescently-tagged cholera toxin subunit B. Retrograde labeling was observed in both primary and higher-order visual areas in hemispheres ipsilateral and contralateral to the injection site (Figure 3A-B). However, the strongest labeling was present in the RSP, a multimodal region that also receives robust input from earlier visual areas ^18,21,24^. In converse experiments, anterograde viral tracing targeting the RSP led to a high density of labeled axons in deeper layers of the dorsal MEC (Supplemental Figure 4).

To determine whether this pathway contributes to MEC visual responses, we expressed the inhibitory opsin eOPN3 ^25^ in the retrosplenial cortex (see Methods). Histological analysis of eOPN3-GFP-expressing fibers again confirmed strong labeling in the deeper layers of dorsal MEC (Figure 3C, Supplemental Figure 4). We recorded MEC activity before and after brief optical stimulation through an implanted fiber (see Methods). Optogenetic suppression of RSP terminals robustly reduced MEC visual response magnitudes that recovered after 30 minutes (baseline SNR: 0.68±0.63; post-illumination SNR: 0.12±0.32SNR; recovery SNR: 0.54±0.50SNR, n= 22 cells in 4 mice, p<0.001 and p=0.003 for illumination and recovery, respectively, linear mixed effects model, Figure 3D-E). Optogenetic suppression did not affect the spontaneous firing rate of MEC neurons (baseline: 1.52±1.35 Hz; post-illumination: 2.03±2.13Hz; recovery: 1.62±1.92 Hz, n= 22 cells in4 mice, p=0.178 and p=0.227, respectively, Figure 3E), and optical stimulation in the absence of eOPN3 expression did not alter MEC activity (Supplemental Figure S4). Overall, these results demonstrate that show that RSP inputs are a dominant pathway carrying visual information to the MEC.

## Discussion

In the present study, we found that single neurons in the MEC robustly respond to simple visual stimuli that are uncoupled from behavioral relevance. Indeed, like other higher order visual areas, responsive MEC cells exhibited contrast sensitivity and large receptive fields but minimal tuning for other stimulus features such as orientation, spatial frequency, and motion. These results suggest that cognitive associations between external visual cues and internal representations of the environment may arise either within or downstream of the MEC.

Visually responsive cells were primarily located within the deeper layers (3 and 5) of the dorsal MEC, a conclusion supported by activity-dependent labeling using both cFos-Trap mice and the activity-dependent photoconversion of the genetically encoded fluorophore CaMPARI2. Individual MEC neurons varied in their representations of stimulus dynamics. Transient cells most sensitive to stimulus onset exhibited significantly higher visual response magnitudes, were more modulated by locomotion, and were more correlated within-group in comparison to sustained cells, suggesting these two broad functional categories may correspond to distinct subpopulations of neurons within the MEC. Indeed, the higher noise correlation values suggest a greater degree of either synaptic interconnectivity or shared inputs ^23^. Ex vivo recordings indicate that recurrent connectivity is considerably stronger in deeper versus superficial layers of the MEC ^26^, suggesting the potential for selective amplification of afferent visual inputs by specific subpopulations. Moreover, approximately 30% of CaMPARI2-labeled cells were positive for the molecular marker Ctip2. It is intriguing to speculate that transient and sustained cells may thus be divided into Ctip2-positive and -negative populations with differing degrees of recurrent synaptic connections that support varying magnitude of visual responses.

The localization of visually-responsive MEC neurons in layers 3 and 5 suggests they may be distinct from other functionally-defined groups including grid and time cells which have largely been described in layer 2 ^27,28^, although speed cells have been described across all MEC layers ^8^. Outputs from MEC layer 3 targets the apical dendrites of pyramidal neurons in the hippocampus, consistent with a functional link between MEC spatially tuned cells and CA1 place cells ^29-32^. Interestingly, silencing MEC output produces an expansion of CA1 place fields ^29-31^. Furthermore, recent work suggests a role for specific environmental cues to trigger input to CA1 that induces behavior timescale dependent plasticity ^33,34^, a function that may be mediated by Layer III visual responding cells.

Our optogenetic results indicate that projections to the MEC from the RSP make a major contribution to entorhinal visually evoked activity, consistent with previous anatomical and ex vivo electrophysiological studies ^18,21,24^. The RSP receives direct sensory inputs from lower-level visual areas ^35-37^, and RSP neurons can encode encode visual landmark information during navigation in a virtual environment ^17^. In addition, cells in the RSP may represent relationships between behaviorally salient environmental cues ^17^, though whether this property arises in the RSP or is inherited from other areas is unclear. As out results show that MEC neurons can encode simple visual stimuli, and the MEC also sends a return projection back to the RSP, it is possible that associations between stimuli and with behaviorally salient cues are formed within the hippocampal formation and then directed back to the RSP.

Visual information is critical for accurate navigation of the environment ^4,9-11^, and visual inputs influence both place and grid cells during exploration ^5,9,11-13^. Similarly, loss of visual information can alter environment-specific activity in both the hippocampus and MEC ^11,12,14^. Computational studies suggest visual signals may serve as a source of error correction during path integration ^9,15,16^. In addition to directly encoding visual stimuli, we also find that visual inputs increase the phase locking of MEC neurons to ongoing theta oscillations. Indeed, theta-band activity is associated with active navigation, coordinating grid cell activity and influencing synaptic plasticity at entorhinal-hippocampal connections ^13^. Thus, visual signals may indirectly regulate a variety of circuit dynamics central to navigation and memory formation.

MEC neurons exhibit a variety of navigation-specific dynamics, variously encoding spatial grids that tile the local environment, regional edges or borders, specific landmarks, task timing, and locomotion speed. As the present work was carried out in animals not actively navigating, it will be extremely interesting to explore whether MEC neurons responsive to visual cues also exhibit coding for these other environmental features or comprise a wholly distinct population. Similarly, the impact of discrete visual stimulus features on other navigation encoding properties of MEC neurons is largely unknown. Future studies taking advantage of high density recordings in mice exploring virtual environments will be critical to addressing the intersection of such representations within the MEC.

## Acknowledgements

The authors thank members of the Higley laboratory for helpful input throughout all stages of this study. This work was supported by funding from the NIH (MH099045, MH121841, MH113852, and EY033975 to MJH, EY026878 to the Yale Vision Core).

## Author Contributions

OD and MJH designed the study. OD and MJH developed the analytical approach. OD carried out all experiments and analyses. OD and MJH wrote the manuscript.

## Competing Interests Statement

The authors declare no conflicts of interest exist.

## Material & Methods

### Animals

C57Bl/6 and FosTRAP heterozygous male mice were kept on a 12h light/dark cycle, provided with food and water ad libitum. All animal handling and experiments were performed according to the ethical guidelines of the Institutional Animal Care and Use Committee of the Yale University School of Medicine.

### Surgical Procedures

Mice were anesthetized with isoflurane (1.5% in oxygen) and maintained at 37°C for the duration of the surgery. Analgesia was provided with subcutaneous injections of Carpofen (5mg/kg) and Buprenorphine (.05mg/kg). Lidocaine (1% in 0.9% NaCl) was injected under the scalp to provide topical analgesia. Eyes were protected from desiccation with ointment (Vetlube). For retroviral tracing of MEC inputs, the scalp was gently cut and a small craniotomy was executed at the coordinates of the left MEC (AP angle -10°, AP: 500µm anterior to the lambdoid suture, LM: -4,2mm), and the injections of Cholera Toxin Subunit B (Recombinant), Alexa Fluor™ 555 Conjugate (ThermoFisher, 200nl measured using a Nano Injector, 2mm and 1.5mm deep from the brain surface) was performed using a thin glass capillary (40-80µm at the tip). After injection, the capillary was maintained in situ for about 10 min before being slowly removed. The scalp was then sutured and the animal housed with its mates for 1 week before perfusion. For acute head fixed recording of the MEC, the scalp was resected and the skull cleaned with Betadine. A surgical screw was implanted on the skull between the eyes and custom headpost were glued to the skull above the bregma suture. Ground and reference wires were inserted above the cerebellum. A circular plastic ring (∼1.5mm diameter) was glued on the skull above the left MEC (centered at AP: 500µm anterior to the lambdoid suture, LM: -4,2mm). All custom head implants were affixed to the skull with dental cement (Metabond, Parkell Industries). For optogenetic manipulations of RSC terminations inside the MEC, the scalp was gently cut and two craniotomies were performed above the left RSC (AP: -4 mm and -3,6mm LM: - 0,6mm), 200nL of AAV9-CaMKII-Cre (10^12 gcp/mL Addgene) was injected at a depth of 300µm in each craniotomies. After 2 weeks the same procedure was used to injected 200nL of AAV5-hSyn1-SIO-eOPN3-mScarlet-WPRE (3,4 10^12 gc/mL Addgene) at the same previous locations inside the RSC, headpost and ground and reference were implanted for later acute electrophysiological recording. For CaMPARI 2.0 tagging of visual responding cells, the scalp was resected and the skull cleaned with Betadine and a small craniotomy was executed at the coordinates of the left MEC (AP angle -10°, AP: 500µm anterior to the lambdoid suture, LM: -4,2mm), and the injections of AAV5-hSyn-NES-his-CaMPARI2-WPRE-SV40 (2.5x10^12gc/ml Addgene 200nl measured using a Nano Injector, 2mm and 1.5mm deep from the brain surface) was performed using a thin glass capillary (40-80µm at the tip). After injection, the capillary was maintained in situ for about 10 min before being slowly removed. Then, a tapered tip optical fiber (DoricLenses) was gently inserted at the same coordinates inside the MEC (2 mm deep) and affixed with head implant to the skull with dental cement. For fos-TRAP visual responding cells tagging, the scalp was resected and the skull cleaned with Betadine and head implant was affixed to the skull with dental cement.

### Electrophysiological Recordings

Mice were habituated to handling and head fixation on a wheel for 3-5 days prior to electrophysiological recordings. One small craniotomies (∼0.4mm apart) was performed above the left MEC under isoflurane anesthesia. Analgesic was provided as described above and mice were moved back for >2h in their home cage to recover from anesthesia. Mice were head-fixed on the wheel. The ring situated above the MEC was filled with artificial cerebrospinal fluid (ACSF; in mM: in mM: 135 NaCl, 5 KCl, 5 HEPES, 1 MgCl2, 1.8 CaCl2 [adjusted to pH 7.3 with NaOH]), and A1x32-Poly3-10mm-25, 32 channels probe (Neuronexus) or a NN1620 optotrod (for optogenetic manipulation – Neuronexus) were covered with DiI (Molecular Probes) and lowered into the MEC (depth 1482.42±186.16µm n=33 mice).

### Visual Stimulation

Visual stimulation and behavior hardware were generated using the Psychtoolbox Matlab extension (Kleiner et al., 2007) and displayed on a 17’’ by 9.5’’ monitor situated 20cm in front of the animal (Binocular experiment) or 15 cm from the right eye (all other recording sessions). Screen display was linearized and maximum luminance was adjusted to ∼140 cd.sr/m2. An iso-luminant grey background was displayed between visual stimuli. Drifting or static Gratings were presented for 1s and separated by a 4s interstimulus interval. Unit responses properties were investigated independently at 4, 20 and 100% contrasts, 0.01, 0.0275, 0.045, 0.0625 and 0.08 cycle/degree spatial frequencies, 5, 10, 40 and 100 size degree and 0, 54, 108, 154, 205, 257, 308 and 360 degree orientation. For optogenetic silencing of RSC terminations inside the MEC using eOPN3, 4s of continuous green pulse (5 mW at the tip MINTF4 - 554 nm, Fiber-Coupled LED Thorlabs) was delivered every minute through the fiber coupled with the probe during visual presentation. In all experiments, pupil size were recorded at 10Hz using an infrared camera (FLIR). Local Field Potentials, wheel motion, and timing signals for face movies, visual stimulus, and behavior were acquired at a 40kHz sampling rate (DigitalLynx system). At the end of the recording session, animals was transcardiacly perfused with 4% paraformaldehyde.

### Activity-dependent labeling of MEC neurons

FosTRAP mice were habituated to handling and head fixation on a fixed wheel for 3-5 days prior light deprivation for 7 days. Tamoxifen was dissolved at 20 mg/ml in pure ethanol, then dissolve at 1/2 in a 1:4 oil mix (Castor oil: sunflower oil) at 37°C for 3 hours. The dissolved tamoxifen was injected immediately intraperitoneally at 20 mg/kg right before mice were positioned on a fixed wheel. Visual presentation were displayed on a 17’’ by 9.5’’ monitor situated 20cm in front of the animal or 15 cm from the right eye in the same condition described before (300 Drifting-gratings patches were presented for 1s and separated by a 4s interstimulus interval with 100% contrasts, 0.04 cycle/degree spatial frequencies, 100 size degree and 0 degree orientation). Animals were then transcardially perfused with 4% paraformaldehyde. CaMPARI 2.0 photoconvertion: mice were habituated to handling and head fixation on a wheel for 3-5 days prior experiments. Mice were head-fixed on the wheel and a 405nm LED (Thorlab) with a FMA-ferrule connector cable (Thorlab) was connected to the implanted cannula and mice . Visual stimulations were presented simultaneously with 405 nm light delivery (2mW at the tip with 300 Drifting-gratings patches were presented for 1s and separated by a 4s interstimulus interval with 100% contrasts, 0.04 cycle/degree spatial frequencies, 100 size degree and 0 degree orientation). Animals were then transcardially perfused with 4% paraformaldehyde in Sorenson’s buffer.

### Histology

Coronal or sagittal sections (50µm-thick, cut using a VT1000S Leica vibratome) were mounted and coverslipped on slides DAPI. Widefield imaging of DiI labeling for post-hoc verification of probe position relative to the MEC or confocal stacking of CTB, eOPN3 and fos expression were acquired (555 nm LED/laser) and visualized with the ImageJ software. For CaMPARI2.0 immunostaining experiments,50 µm-thick slices were washed 2 times with a solution of 0.3% Triton X-100, 10 minutes each and then incubate at room temperature during 4-6 hours. Primary antibodies: mouse anti-Campari red (1:500 Absolute Antibody) and Guinea-Pig anti Ctip2 (1:500 Synaptic System) or rabbit anti-reelin (1:500 invitrogene) or rabbit anti-calbindin (1:500 Synaptic System) were diluted in blocking solution (0.02 % BSA, 10% goat serum, 0.5% Triton X-100) and slices were incubated overnight at 4°C on a rocking table. Slices were rinsed 4 times with 300ul blocking solution (15 minutes each) and incubated with secondary antibodies diluted in blocking solution (goat anti-mouse conjugated with AlexaFluor 555 (ThermoFisher) and goat anti-guinea-pig conjugated with AlexFluor 647 or goat anti-rabbit conjugated with AlexaFluor 647) for 1 hour at room temperature. Slices were then rinsed 4 times, then mounted and coverslipped with DAPI. Slides were allowed to dry overnight before imaging with 385 nm, 488 nm, 543 nm, and 647 nm lasers on a confocal microscope.

### Data Analysis

Electrophysiology data were analyzed in Matlab 2020b (Mathworks) using custom scripts. All time-series were down-sampled to 20KHz and aligned. Pupil diameter was computed from movies using FaceMap (Stringer et al., 2019). First, data were Z-score normalized within a session, and the high and low thresholds corresponding to 60% and 40% quantiles in the data distribution, were extracted for that session. Data were then smoothed using a 1-s window moving-average filter, and epochs during which smoothed data continuously exceeded the high and low Z-thresholds for at least 5 s were considered high and low Pupil size, respectively. Wheel position was determined from the output of the linear angle detector. The circular wheel position variable was first transformed to the [−π, π] interval. The phases were then circularly unwrapped to obtain running distance as a linear variable, and locomotion speed was computed as a differential of distance (cm s −1). A change-point detection algorithm detected locomotion onset and offset times based on changes in standard deviation of speed for at least 5s to be considered as locomotion periods. Single units were extracted and clustered from LFP recording kilosort2. Resulting clusters were manually verified and refined using the phy-gui (https://github.com/cortex-lab/phy). Pupil diameter and locomotion were interpolated and aligned to the spike time data.

### Identification of visual responding cells

Individual neurons activity post-stimulus time histogram (PSTH peri-windows size 2000 ms and bins size 10 ms) were computed at the onset of the same visual stimulus (1s duration every 4s, drifting grating, 100% contrast, 0° Orientation, 100° size, 0.04 Hz spatial frequency, 2Hz temporal frequency). Same PSTH ware computed by shuffling the inter spikes interval 1000 times for each neurons giving an average shuffling PSTH. Neurons were considered as positively responding if the mean of the raw PSTH was greater than the mean of the shuffling PSTH during the visual presentation (p value>0.01). cells were categorized as inhibited responding cells when the mean of the raw PSTH was significantly lower than the mean shuffling PSTH (p value>0.01). Signal noise ratio of neuronal visual response was calculated as Δ/baseline where Δ is the difference between the mean firing rate during the stimulation and the mean firing rate in baseline.

### Behavioral state modulation of neural activity

Dynamics of locomotion and pupil size were calculated with a modulation index: (FRwin-FRbaseline)/(FRwin+FRbaseline) where FRwin is the neuron firing rate during locomotion and large pupil size and where FRbaseline is the neuron firing rate during immobility and small pupil size. Modulation of visual response by ongoing behavior was calculated with the same metric: (SNRwin-SNRbaseline)/(SNRwin+SNRbaseline) where SNRwin is the SNR during visual presentation during locomotion and large pupil size and where SNRbaseline is the SNR during visual stimulation during immobility and small pupil size. Because Large Pupil size is highly correlated with locomotion, large Pupil size periods during locomotion were exclude from these analysis. Noise correlation were performed for all pairs of neurons for each groups (described as quartiles from the ratio A/B distribution) using the Pearson correlation coefficient.

### Theta modulation of neural activity

Identification of theta periods: the raw LFP was down sample to 1250 Hz and integrated power was calculated in the 4-11Hz (theta) and 1-3Hz (delta) frequency bands using the multi-taper method (1 s time window and 4 tapers) implemented in the Chronux MATLAB toolbox ^38^. Theta periods were defined as theta/delta power ratio above 3 and validated manually using the Sonic Visualizer free software. Theta phase was computed in theta periods using the Hilbert transform on band-pass filtered (4-11 Hz) down-sampled LFP. Phase modulation of neurons relative to theta oscillations was tested by using the Rayleigh test (nonuniformity of the circular distribution). When phase modulation was significant (p < 0.05), von Mises parameters µ and κ were estimated via maximum likelihood as indices of preferred phase and modulation strength, respectively (CricStat MATLAB tool box (Berens, 2009 #1693)). The number of CTB or td-tomato expressing cell bodies, were calculated with Imagej object counter plugin and processed with the SHARCQ matlab GUI ^39^ for atlas alignment. CaMPARI2.0 colocalization and fluorescence intensity where measured with CellProfiler plugins ^40^. Briefly, we identified the location of the tapered-tip optical fiber relative to the CaMPARI2.0 infection for calculating the green-to-red photoconversion. We detected the cell bodies in the 488 nm and 555 nm channels to identify the photoconverted cells. We then quantified the colocalization of red cells with the secondary antibodies with the 647 nm channels to quantify the amount of cells expressing the targeted molecular markers.

### Statistics

All statistical analyses were conducted using custom-written scripts in MATLAB. We used mixed effect regression models for electrophysiology data, due to its nested structure with multiple cells recorded within each mouse. We treated the experiment type the fixed effect, and the individuals (mice) were random effects. Since our experimental design was between-subject, we used the fitlme function in MATLAB to fit the intercepts of the random effects, with the response modeled as: response ∼ experimentType + (1 | mouse). For the histological data, given one measurement per mouse, data was compared with either a t-test after a test for normality or an ANOVA test. Each experiment was conducted in one or two recordings per animal. Experiments were not independently replicated. Only animals with sufficient amount of theta, locomotion and large Pupil size periods were included for behavioral analysis. Differences were considered statistically significant at p < 0.05. Unless indicated otherwise, all results are presented as mean ± SD.

---

**Supplemental Figure 1.**
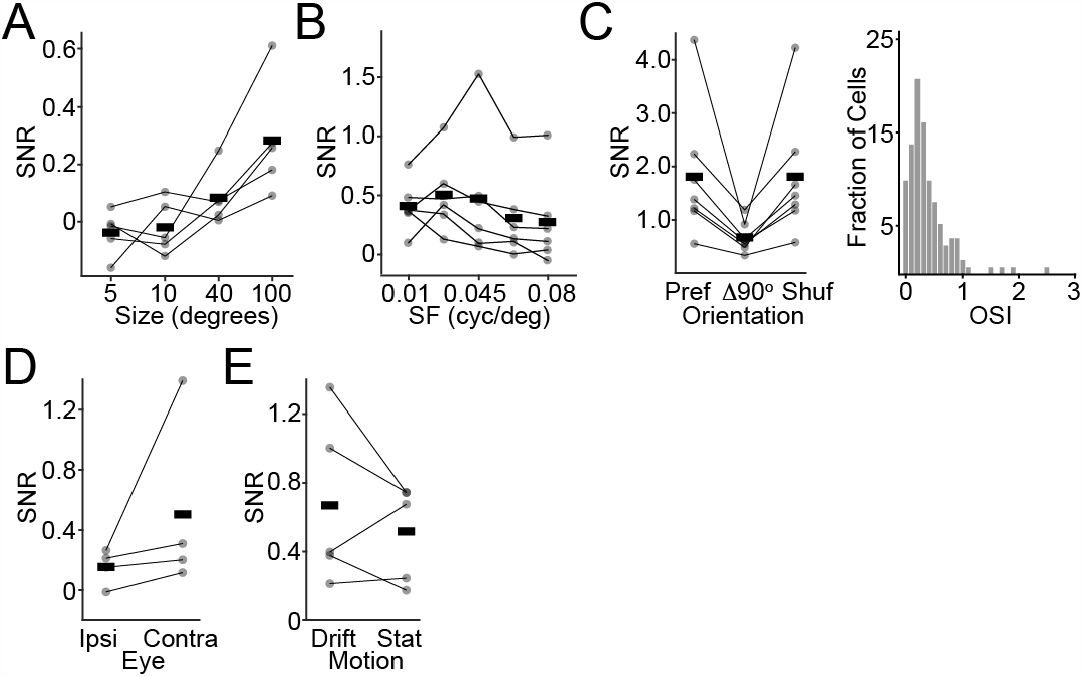
Sensitivity of MEC neurons to visual stimulus properties. **A**, Visual SNR as a function of stimulus size. **B**, Visual SNR as a function of stimulus spatial frequency. **C**, Visual SNR as a function of orientation, showing preferred, 90-degrees orthogonal to preferred, and shuffled controls values (left). Histogram of orientation selectivity indices (right). **D**, Visual SNR for stimulation of the eye ipsilateral or contralateral to the recorded MEC. **E**, Visual SNR for drifting versus static inverting gratings.

**Supplemental Figure 2.**
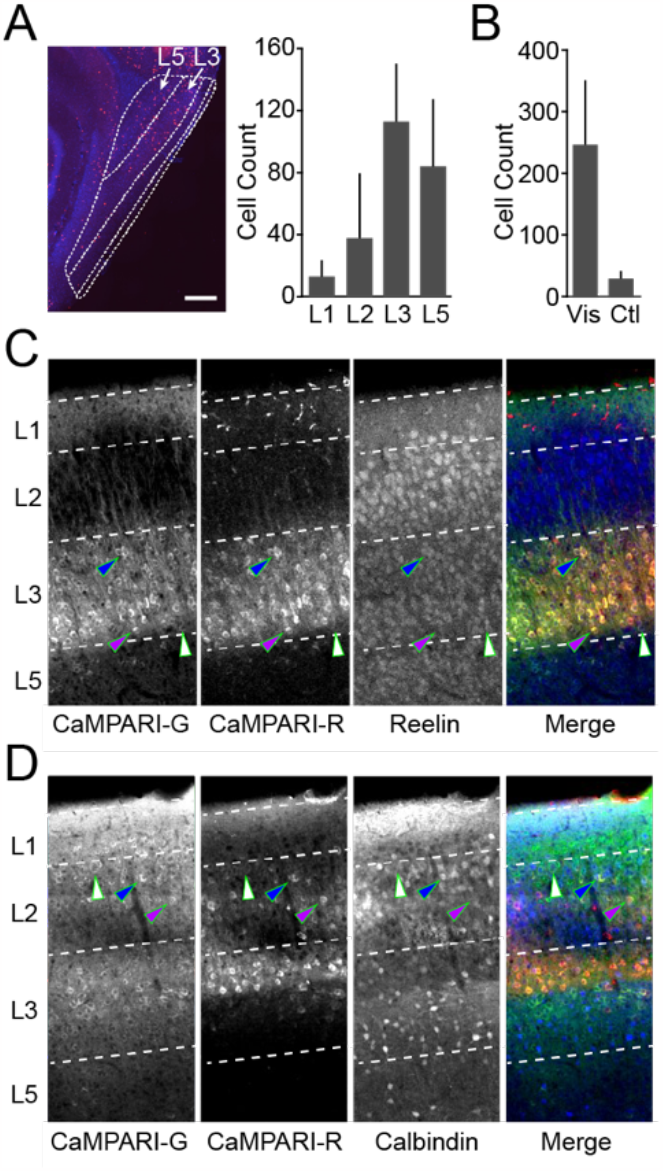
Histological characterization of visually responsive MEC neurons. **A**, Example image showing fos-Trap-labeled visually responsive cells in MEC (left) and average count of fos-Trap labeled cells across MEC layers (right). **B**, Average count of fos-Trap-labeled cells in the MEC for visual stimulation and control (no stimulation) animals. **C**, Example images showing green fluorescent CaMPARI2-labeled cells, photoconverted red fluorescent CaMPARI2-labeled cells, Reelin-positive cells, and merge. Non-converted cell (green arrow), converted reelin-positive cell (blue arrow), and converted reelin-negative cell (purple arrow) are indicated. **D**, As in (C) for calbindin-expressing cells.

**Supplemental Figure 3.**
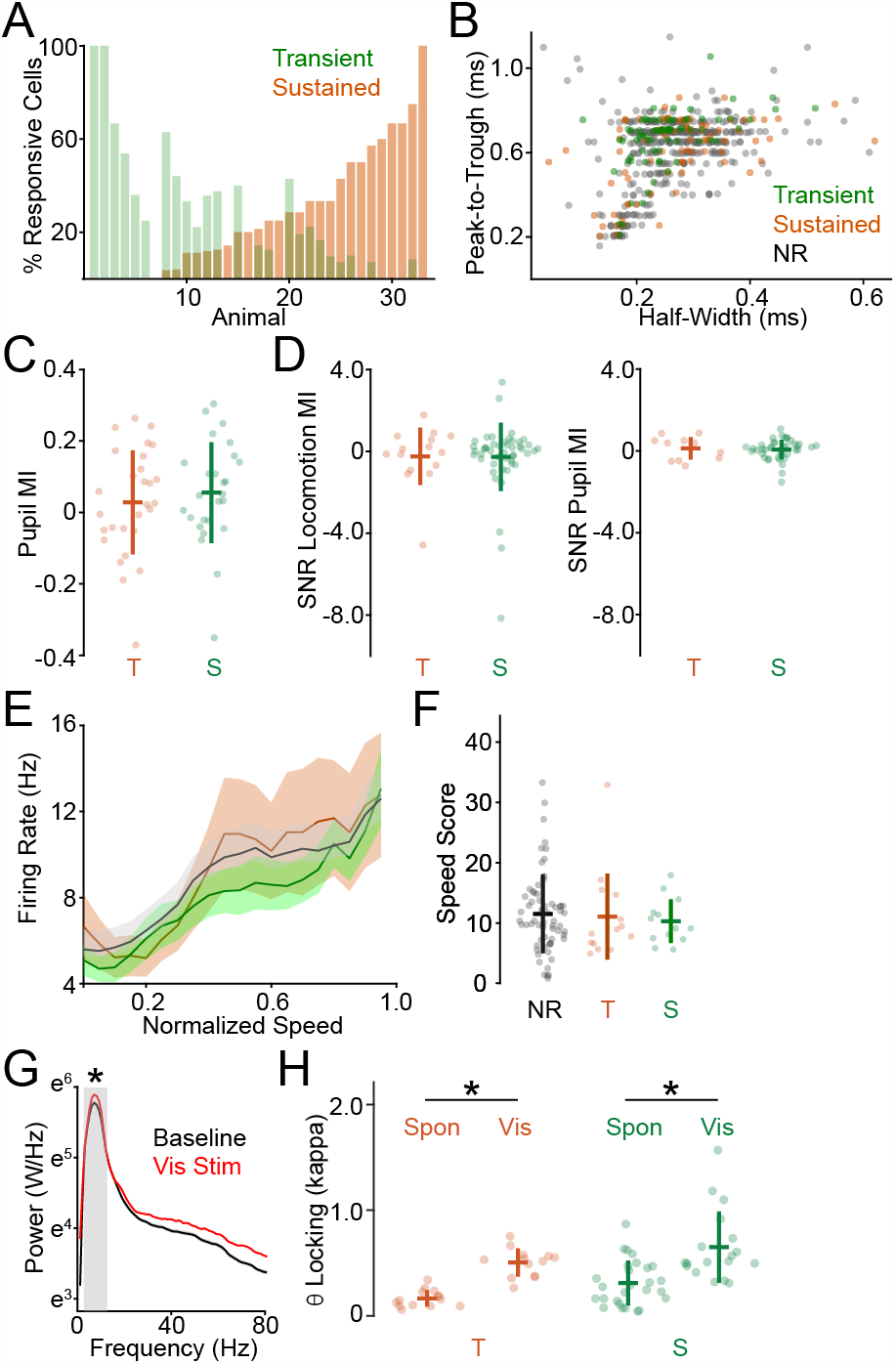
Functional properties of visually responsive MEC neurons. **A**, Fraction of transient versus sustained responsive cells across all animals. **B**, Spike waveform features for all neurons across all animals, with colors indicating transient, sustained, or non-responsive neurons. **C**, Pupil modulation index for spontaneous activity in transient versus sustained cells. **D**, SNR Modulation Index for locomotion (left) and pupil diameter (right) for transient versus sustained cells. **E**, Average spontaneous firing rate as a function of running speed for transient versus sustained cells. **F**, Average speed score for non-responsive, transient, and sustained cells. **G**, Average power spectrum density across animals for spontaneous activity and during visual stimulation. Gray box indicates theta range. **H**, Average theta-locking for single units for spontaneous activity and during visual stimulation, for transient versus sustained cells. * indicates p<0.o5 (see Main Text).

**Supplemental Figure 4.**
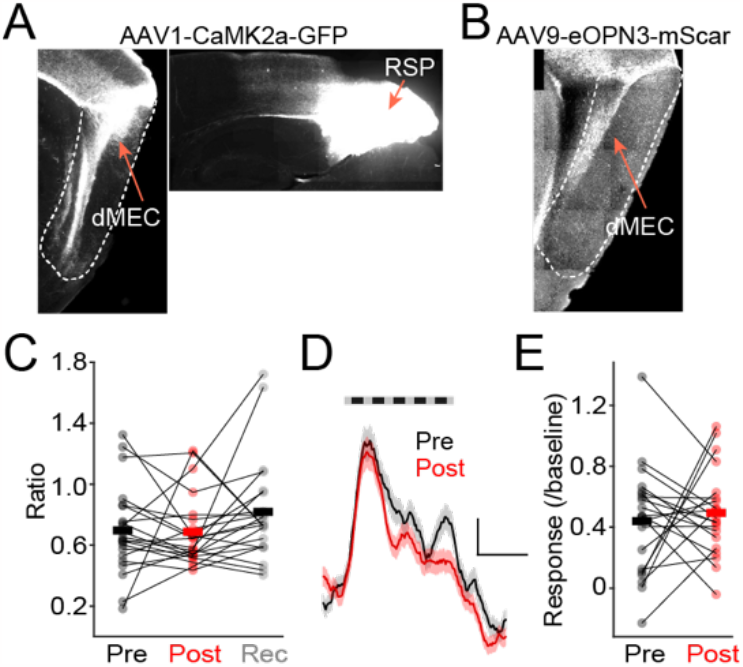
Optogenetic suppression of RSP inputs in the MEC. **A**, Example image showing anterograde labeling of RSP inputs to MEC using AAV-CaMKIIa-GFP tracing. **B**, Example image showing eOPN3-mScarlet-labeled fibers in the MEC originating from RSP. **C**, Control data showing visual response relative to baseline before and after light stimulation in eOPN3-non-expressing animals. **D**, Average visual responses before and after light stimulation in control animals. **E**, Average spontaneous firing rate before and after light stimulation in control animals.

## Notes

### Competing Interest Statement

The authors have declared no competing interest.

